# A putative xanthine dehydrogenase protects *Borrelia burgdorferi* from reactive oxygen species and promotes virulence

**DOI:** 10.1101/2022.08.24.505212

**Authors:** James P. Phelan, Jeffrey S. Bourgeois, Julie E McCarthy, Linden T. Hu

## Abstract

*Borrelia burgdorferi* is a pathogenic bacterium and the causative agent of Lyme Disease. It is exposed to reactive oxygen species (ROS) in both the vertebrate and tick hosts. While some mechanisms by which *B. burgdorferi* ameliorates the effects of ROS exposure have been studied, there are likely many other unknown mechanisms of ROS neutralization that contribute to virulence. Here, we follow up on a three gene cluster of unknown function, *BB_0554, BB_0555*, and *BB_0556*, that our prior unbiased transposon insertional sequencing studies implicated in both ROS survival and survival in *Ixodes scapularis*. We confirmed these findings through genetic knockout and provide evidence that these genes are co-transcribed as an operon to produce a xanthine dehydrogenase. In agreement with these results, we found that *B. burgdorferi* exposure to either uric acid (a downstream product of xanthine dehydrogenase) or allopurinol (an inhibitor of xanthine dehydrogenase) could modulate sensitivity to ROS in a *BB_0554-BB_0556* dependent manner. Together, this study identifies a previously uncharacterized three gene operon in *B. burgdorferi* as a putative xanthine dehydrogenase critical for virulence.

**Importance:** *Borrelia burgdorferi*, the causative agent of Lyme disease, is highly successful at evading host immune defenses such as reactive oxygen species, despite minimal identified defenses against oxidative stress. We identified a putative xanthine dehydrogenase that is important in survival of the organism when exposed to oxidative stress and in both its tick and murine hosts. The mechanism appears based on a previously unrecognized role of uric acid in neutralizing reactive oxygen species and highlights how *B. burgdorferi* can utilize its very limited metabolic capabilities in unique ways.

## Introduction

*Borrelia burgdorferi* is the causative agent of Lyme disease, the most common arthropod borne disease in the United States (1). The spirochete is maintained through an enzootic cycle that includes *Ixodes* ticks and wide array of mammalian and avian reservoir hosts (1), and thus must be able to identify and respond to stressors from these vastly different environments. One such stressor seen throughout the enzootic cycle is reactive oxygen species (ROS). After *Ixodes* ticks take a bloodmeal, ROS are detectable within the salivary glands as well as the midgut (2,3). Additional ROS are also likely encountered by the bacterium immediately after entering a mammalian host, as recruited immune cells attempt to leverage these species to destroy the invading pathogen (4,5). These reactive oxygen species can cause widespread damage to *B. burgdorferi* membranes (6), and yet during successful infections they do not—suggesting that the bacteria has evolved mechanisms to overcome this early barrier.

Interestingly *B. burgdorferi* appears capable of surviving ROS insult despite encoding a very limited selection of ROS detoxifying proteins, particularly compared to the canonical ROS-detoxifying pathways of *E. coli*. While a small number of genes (including a manganese transporter (BmtA), a manganese-dependent superoxide dismutase (SodA) (7–9), and a Coenzyme A reductase (10,11) have been implicated in *B. burgdorferi* resistance to ROS, it remains likely that additional pathways exist to aid survival. Our lab previously utilized a transposon insertional sequencing (Tn-seq) screen to identify additional genes that are involved in *B. burgdorferi*’s response to ROS (12). That work confirmed the importance of three transmembrane proteins (BB_0017, BB_0164, and BB_0202) in contributing to ROS resistance and virulence, however, there were additional genes identified by the screen that could play important roles in *B. burgdorferi* pathogenesis.

In this work we attempted to follow-up on candidates from this prior screen in order to identify ROS-resistance genes that are critical for *B. burgdorferi* to survive ROS defenses during transmission. To do this, we leveraged data from a second published Tn-seq screen, which identified genes required for survival in the *Ixodes scapularis* larvae following feeding (13). By examining genes that appear in both screens, we identified the *BB_0554, BB_0555*, and *BB_0556* locus as critical for ROS resistance and survival following the tick bloodmeal. Here, we tested the importance of these genes through genetic knockout and complementation, confirming that they play a key role in ROS resistance. Interestingly, we also note the operon is required for murine infection independent of the tick, demonstrating a key role for the operon in multiple stages of infection. Finally, to understand the function of this locus, we utilized *in silico* and experimental tools to identify a putative role for the locus as a three-subunit xanthine dehydrogenase. These results suggest that targeting xanthine metabolism could help to block *B. burgdorferi* transmission.

## Results

### Integration of two Tn-seq screens to identify genes involved in ROS survival during the tick bloodmeal

During the tick bloodmeal, *Borrelia burgdorferi* is exposed to a plethora of nutrients, metabolites, and signaling molecules. While some components of this bloodmeal are useful for the bacteria, reactive oxygen species (ROS) are also present and represent one of the first hurdles *B. burgdorferi* must overcome to successfully colonize a mammalian host. We hypothesized that we could leverage two previous transposon insertional sequencing (Tn-seq) screens—one that screened for ROS resistance *in vitro* (12) and one that screened for survival in *Ixodes scapularis* larvae following a bloodmeal (13)—to identify *B. burgdorferi* genes involved in surviving this early selective pressure. To do this, we examined transposon mutants in which (a) there was a 50% reduction in relative frequency within the pool after exposure to H_2_O_2_ compared to input (12), and (b) no mutant was recovered after larval tick repletion despite robust presence in the input library (13) **(Figure 1A)**. Using these criteria, we identified 9 genes that were shared between the two screens **(Figure 1B, Table 1)**. Among this list, we identified a two gene locus comprised of *BB_0554* and *BB_0555* that is important for ROS defense and survival in larval ticks. Notably, we observed a third gene in this region, *BB_0556* in this locus that barely missed the selection cutoff but showed survival trends similar to *BB_0554* and *BB_0555* disruption **(Figure 1C)**. Disruption of any of the three genes also rendered the bacteria sensitive to TBHP-induced ROS damage **(Figure 1C)**.

**Table 1:** Genes required for both H_2_O_2_ resistance and survival in tick larvae following bloodmeal.

| Locus | H<sub>2</sub>O<sub>2</sub> Resistance (Frequency Ratio*) | Survival in *Ixodes scapularis* (Frequency Ratio*) |
| --- | --- | --- |
| BB_0017 | 0.256 | 0.000 |
| BB_0042 | 0.414 | 0.006 |
| BB_0411 | 0.313 | 0.006 |
| BB_0412 | 0.249 | 0.000 |
| BB_0420 | 0.417 | 0.000 |
| BB_0554 | 0.312 | 0.003 |
| BB_0555 | 0.352 | 0.001 |
| BB_0597 | 0.438 | 0.013 |

\*Frequency Ratio: Frequency of Tn mutants following selection/Frequency of Tn mutants in input pool. For Tn mutants where no mutant was recovered following selection, value was changed from “0” to “1” for calculation. Ratio represents the average frequency ratio of all mutants containing an insertion event within the listed gene.

**Figure 1:**
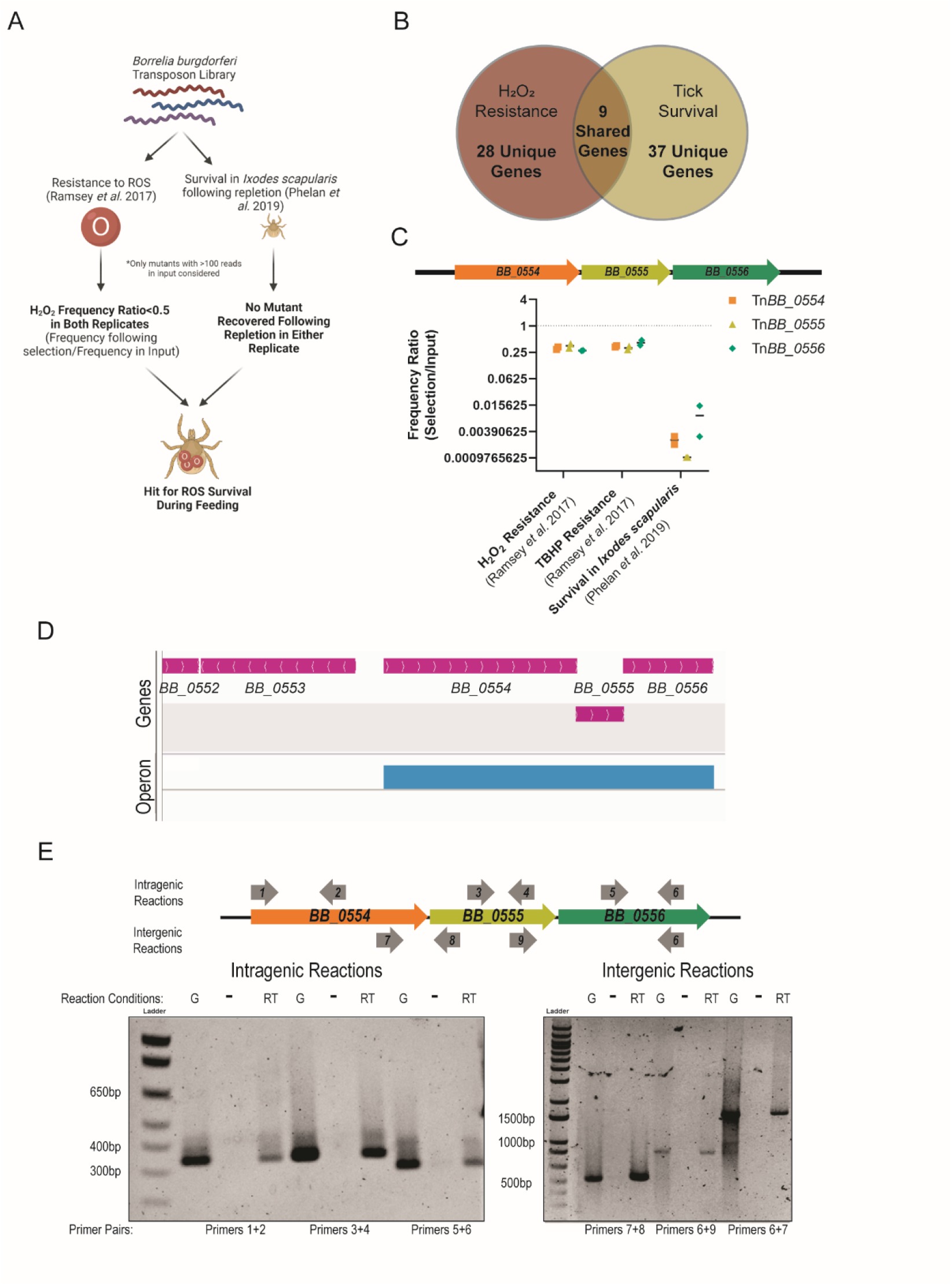
Overlap between two transposon screens identifies a potential role for the *BB_0554-BB_0555-BB_0556* operon in ROS resistance and tick survival. (A) Schematic of criteria used to identify overlap between Ramsey *et al*. (12) and *Phelan et al*. (13) screens. Created with Biorender.com. (B) Venn diagram demonstrating overlap of this from the ROS and *Ixodes* screens. (C) Transposon insertion into *BB_0554, BB_0555*, or *BB_0556* diminishes *B. burgdorferi* fitness following ROS insult or in larval ticks following bloodmeal. Genetic locus is displayed graphically with each arrow representing a gene and the direction representing the direction of transcription. Frequency ratio represents the frequency of the transposon mutants following selection divided by the frequency of the mutants measured in the input pool. Frequency ratio is the average of all transposon insertions within each gene. For insertions where no mutants were recovered following selection, the number of recovered mutants was set to “1” rather than “0” to enable the calculation. Each dot represents an independent Tn-seq experiment and bars represent the mean. (D) Rockhopper software (17) predicts that the *BB_0554-BB_0556* locus is transcribed as an operon. Panel adapted from the IGV genome browser visualization (18). First track shows annotated genes in the *B. burgdorferi* B31 genome with strand denoted by arrow direction. The second track denotes predicted operons based on RNA-seq data (14). (E) RT-PCR confirms *BB_0554-BB_0556* is expressed as an operon. Primer pairs were designed as shown on the schematic in order to amplify genomic DNA (“G”) or reverse transcribed RNA (RT). A no reverse transcriptase RNA control (-) was included to confirm there was no contaminating genomic DNA in the reverse transcribed samples. All intragenic and intergenic primer pairs successfully produced the expected products across the locus.

Based on the apparent ability for all three genes to phenocopy one another, we hypothesized that they may work in the same genetic pathway. In line with this hypothesis, analysis of previously published transcriptomic data (14) revealed that the region is likely expressed as an operon **(Figure 1D)**. A secondary analysis (MicrobesOnline Operon Predictor) which considers conservation of gene cluster, shared functional category, and intragenic distance to determine the likelihood that the genes are organized into an operon (15,16) also predicted that *BB_0554, BB_0555*, and *BB_0556* are organized in an operon. As final confirmation, we showed using RT-PCR that the three genes are transcribed as a single RNA transcript **(Figure 1E)**. Thus, *BB_0554, BB_0555*, and *BB_0556* make up a transcriptional operon and all appear important to both ROS resistance and survival in fed larval ticks.

### Generation of Δ*BB_0554-BB_0556* and complementation strains

To confirm our findings with the transposon mutants and further characterize the operon, we generated a deletion mutant of the entire three gene locus (Δ*BB_0554-BB_0556*). The deletion mutant was generated by allelic exchange utilizing a linearized plasmid pJP01, which replaced the locus with a streptomycin/spectinomycin resistance allele **(Figure 2A, 2B)**. Observations made with the *ΔBB_0554-BB_*0556 mutant were compared with results from a transposon mutant (transposition occurred in *BB_0556*) used in the original screen to determine consistency across the genetic systems. To confirm the effects of the mutation were specific to the gene locus, we complemented the Δ*BB_0554-BB_0556* strain *in trans* **(Figure 2A, 2B)**. Each strain was frequently observed under darkfield microscopy and confirmed to lack any major morphological or motility defects. Additionally, we did not observe a significant difference in the growth rates of any strain used in our study **(Figure 2C)**. Finally, plasmid content of strains was frequently reconfirmed, and all strains contained comparable plasmid content.

**Figure 2:**
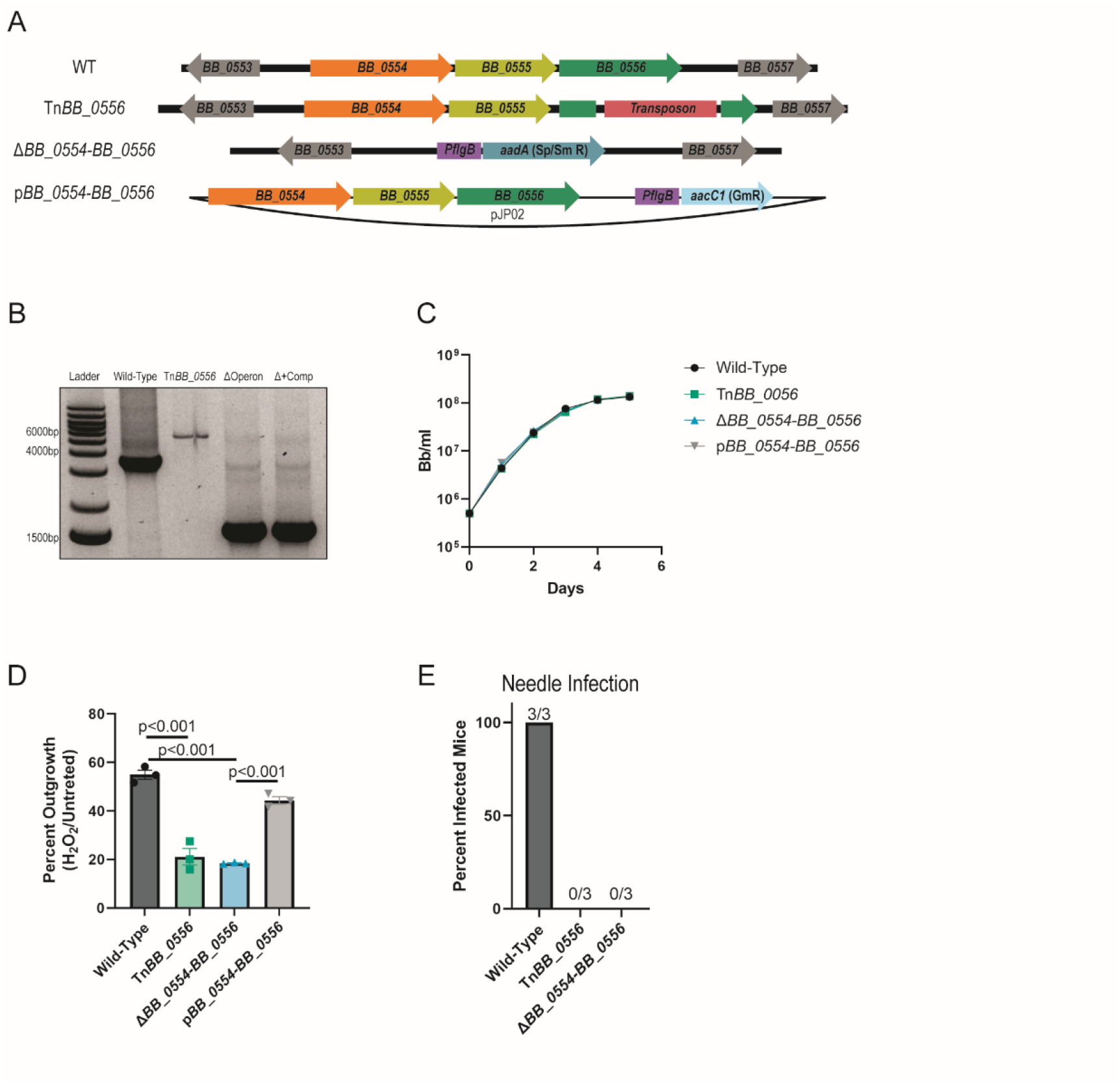
The *BB_0554-BB_0555-BB_0556* operon contributes to ROS resistance and virulence. (A) Schematic of the *BB_0554, BB_0555*, and *BB_0556* locus in strains used in this study. (B) PCR size confirmation of the *BB_0554, BB_0555*, and *BB_0556* locus used in this study. PCR utilized flanking primers downstream of *BB_0553* (primer OperonScreenF) and in *BB_0557* (primer OperonScreenR). Note this reverse primer targets a region outside of the complementation plasmid, and thus only the endogenous locus is amplified. (C) The *BB_0554, BB_0555*, and *BB_0556* locus is not required for growth in BSK-II media. Strains were grown to late exponential phase and diluted to 5×10^5^ *B. burgdorferi* (Bb)/ml. Bacteria were counted by dark field microscopy every 24 hours using a Petroff-Hauser chamber. Each strain was counted in triplicate. Dots represent the mean and error bars represent the standard error of the mean. (D) The Δ*BB_0554-BB_0556* strain recapitulates the ROS sensitivity seen in the Tn*BB_0556* strain and complementation restores resistance. Log phase bacteria were washed and resuspended in modified BSK-II media lacking pyruvate and exposed to 185 μM H_2_O_2_ for four hours at 32°C then subcultured 1:10 in BSK-II. Outgrown bacteria were counted 48 hours later. Data are plotted as 100*(number of outgrown *B. burgdorferi* treated with 185μM H_2_O_2_^/^number of outgrown untreated *B. burgdorferi*). (E) The *BB_0554-BB_0556* locus is required for murine colonization independent of the tick. Subcutaneous injection of either Tn*BB_0556* or Δ*BB_0554-BB_0556* bacteria did not result in detectible colonization of C57BL/6J ear, bladder, or ankles. Tissues were harvested 2 weeks post infection, incubated in BSK-II at 37°C, and monitored for 4 weeks for the presence of spirochetes. Each dot represents one of three independent replicates from a single representative experiment, the bar represents the average, the error bars the independent error of the mean. P-values are from a one-way ANOVA on the log transformed data with Sidak’s multiple comparison test.

### The *BB_0554-BB_0556* locus protects *B. burgdorferi* against reactive oxygen species and promotes virulence

We first determined the ROS sensitivity of each of the newly created strains. We exposed wild-type, tn*BB_0556*, Δ*BB_0554-BB_0556*, and complemented bacteria to H_2_O_2_ and measured their resistance to the stress. Similar to the Tn-seq results, we found that tn*BB_0556* and Δ*BB_0554-BB_0556* bacteria were significantly more susceptible to ROS compared to wild-type or complemented strains **(Figure 2D)**. While the previously published Tn-seq screen demonstrated the importance of the *BB_0554-BB_0556* locus in larval tick survival (13), our focus here was whether the operon was required for overcoming ROS during transmission to the vertebrate host. To test this, we infected C57BL/6J mice with *B. burgdorferi* subcutaneously. Interestingly, we could not recover Tn*BB_0556* nor Δ*BB_0554-BB_0556* bacteria from needle injected mice **(Figure 2E)**. This suggests that *BB_0554, BB_0555*, and *BB_0556* are required for virulence.

### *BB_0554, BB_0555*, and *BB_0556* are predicted to form a xanthine dehydrogenase

Having established the importance of *BB_0554, BB_0555*, and *BB_0556* in *B. burgdorferi* fitness, we next attempted to understand the function of the gene products. Utilizing the NCBI BLAST toolkit revealed that each gene is at least 90% identical among 34 diverse *B. burgdorferi* sensu lato isolates, with *BB_0554* and *BB_0556* nearing 100% identity **(Figure 3A)**. This conservation suggests a critical role for the genes, in line with our observations that they are critical for fitness during the enzootic lifecycle. To gain insight into what conserved role these genes play, we leveraged *in silico* techniques to identify putative functional protein domains in each gene. We first examined *BB_*0554, a putative 622 amino acid protein that is annotated as a hypothetical protein. Utilizing protein prediction software (Phyre2), we found *BB_0554* secondary structure is predicted to have a molybdopterin binding domain with 82% confidence and 12% coverage. Next, *BB_*0555, is a putative 155 amino acid protein that is annotated as a hypothetical protein. Phyre2 structural modeling of this protein indicated a highly conserved iron sulfur subunit found in many different oxidoreductases and dehydrogenases. The confidence of the top ten predicted models ranges from 88 to 95 % with at least 55% coverage. Finally, *BB_*0556 is a 289 amino acid protein that is annotated as a xanthine dehydrogenase (XDH) family protein subunit M. Phyre2 protein prediction indicated a strong homology to many FAD binding subunits of oxidases and dehydrogenases, with models reaching 100% confidence with 98% coverage of the protein.

**Figure 3:**
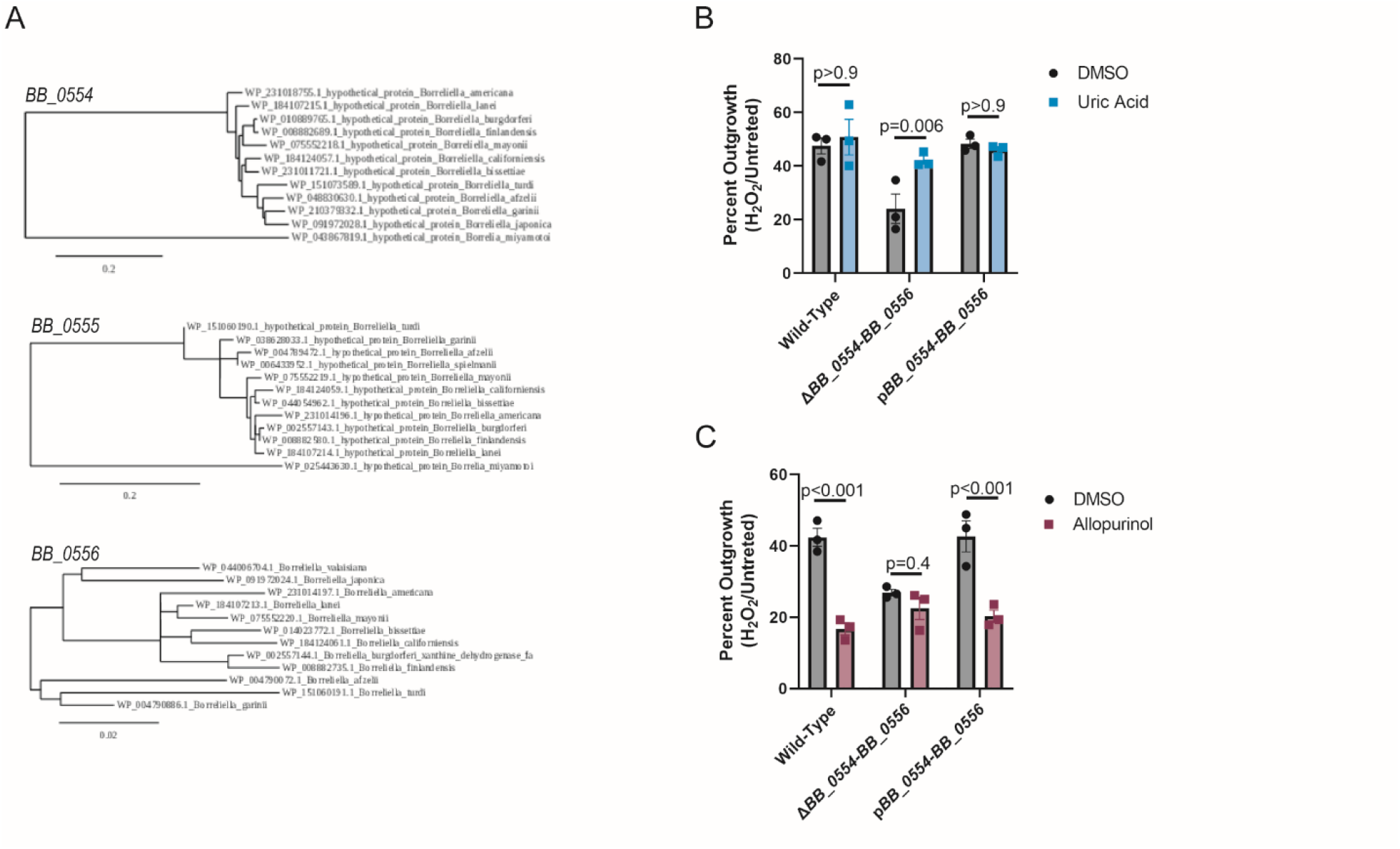
The *BB_0554-BB_0555-BB_0556* operon encodes a putative xanthine dehydrogenase. (A) Phylogenetic analysis of the individual predicted protein sequences sequence among *Borrelia* species. Protein Blast was performed to look for sequence similarities and then each were compared using we service Phylogeny.fr. Distances were estimated using PhyML matrix and is a measure of changes per site. Accession numbers are listed with strain name. (B) Uric acid restores ROS resistance to the *ΔBB_0554-BB_0556* mutant. Log phase *B. burgdorferi* were treated with 300nM uric acid for two hours, then washed and resuspended in modified BSK-II media lacking pyruvate. Bacteria were then exposed to 185μM H_2_O_2_ for four hours at 32°C then subcultured 1:10 in BSK-II. Outgrown bacteria were counted 48 hours later. (C) A xanthine dehydrogenase inhibitor reduces *B. burgdorferi* ROS resistance in a *BB_0554-BB_0556* dependent manner. Log phase *B. burgdorferi* were treated with 1mM allopurinol for two hours, then washed and resuspended in modified BSK-II media lacking pyruvate. Bacteria were then exposed to 0μM or 125μM H_2_O_2_ for four hours at 32°C then subcultured 1:10 in BSK-II. Outgrown bacteria were counted 48 hours later. For B and C, data are plotted as 100*(number of outgrown *B. burgdorferi* treated with H_2_O_2_^/^number of outgrown untreated *B. burgdorferi*). Each dot represents an independent replicate from a single representative experiment. Bars represent the mean, error bars the standard error of the mean. P-values from two-way ANOVA on log-transformed data with Sidak’s multiple comparison test.

Of note, prokaryotic XDH can vary in the subunit composition (19,20). However, the complex depends on the combined enzyme subunits containing domains capable of binding Molybdenum cofactor (Moco) (the predicted BB_0554 function), 2Fe-2S (the predicted BB_0555 function), and FAD (the predicted BB_0556). Additionally, many bacterial species produce xanthine dehydrogenases via genes arranged in operons (19,21,22), comparable with our locus here. Thus, our *in silico* analysis supports a hypothesis that these proteins form a functional xanthine dehydrogenase enzyme.

### Uric acid rescues Δ*BB_0554-BB_0556* ROS resistance

Xanthine dehydrogenases are involved in the breakdown of hypoxanthine to xanthine and xanthine to uric acid. Uric acid is a known scavenger of free radicals and can therefore act as an antioxidant (23–25). We attempted to determine the role of BB_0554-0555-0556, on uric acid production through commercial colorimetric assays (Sigma-Aldrich MAK077), NMR, and mass spectrometry to measure the abundance of uric acid in *B. burgdorferi* lysates. However, the sensitivity of each of these assays proved to be too low to detect levels in *B. burgdorferi*. While we were unable to determine whether the *BB_0554-BB_0556* operon impacts uric acid abundance directly, we tested the functional effects of xanthine metabolism on *B. burgdorferi* ROS resistance using two alternative methods: exogenously manipulating the uric acid pool and blocking xanthine dehydrogenase function.

We hypothesized that if Δ*BB_0054-BB_0056* increases ROS sensitivity due to a reduced intracellular uric acid pool, exogenously supplementing that uric acid pool should restore resistance. Indeed, treating Δ*BB_0054-BB_0056* with uric acid prior to H_2_O_2_ exposure eliminated the enhanced sensitivity of the mutant to ROS compared to wild-type and complemented bacteria **(Figure 3B)**. The addition of uric acid had no impact on wild-type and complemented *B. burgdorferi* ROS sensitivity. Thus, it appears that uric acid-mediated protection is saturated in wild-type bacteria but can be increased further in the mutant—in line with the mutant having a reduced uric acid pool.

### A xanthine dehydrogenase inhibitor increases *B. burgdorferi* ROS sensitivity through *BB_0554, BB_0555*, and *BB_0556*

A second method we utilized to link *BB_0554, BB_0555*, and *BB_0556* to uric acid production was to determine whether the phenotype of the deletion mutant could be mimicked by a xanthine dehydrogenase inhibitor, allopurinol (19,20). We pre-incubated the *B. burgdorferi* with 1mM allopurinol or vehicle control and exposed them to 125μM H_2_O_2_. This concentration of allopurinol was chosen as we did not observe any impact of allopurinol on growth and others have reported effects of allopurinol in the 1-2mM range *in vitro* (26). We found that wild-type and complemented *B. burgdorferi* treated with allopurinol were more sensitive to the same amount of ROS than their untreated controls (**Figure 3C**; p<0.001). The degree to which allopurinol treatment increased sensitivity almost exactly phenocopied the level of sensitivity we see in our *BB_0554-BB_0556* operon mutants. Further, the addition of allopurinol to the Δ*BB_0554-BB_0556* strain did not further decrease resistance **(Figure 3C)**, suggesting that the effects of allopurinol and locus ablation on ROS resistance occurs through the same pathway. Together, these results strongly suggest that the operon encodes a xanthine dehydrogenase which aids in ROS detoxification.

## Discussion

In order to successfully complete its enzootic cycle, *B. burgdorferi* must survive reactive oxygen stress across multiple diverse environments. Here we describe a novel locus that is critical for *B. burgdorferi* to survive ROS, and potentially other insults, that it faces as it begins its journey from the tick vector into a mammal and back. Based on *in silico* and experimental data, we hypothesize that *BB_0554, BB_0555, and BB_0556* encode a xanthine dehydrogenase. Interestingly, prior studies have found that there is a significant decrease in *BB_0554* and *BB_0555* transcripts as *B. burgdorferi* grows from early exponential to stationary phase (14) which is consistent with a role in processing hypoxanthine as there is less need to utilize hypoxanthine and generate uric acid as growth slows.

Our data demonstrate that this putative xanthine dehydrogenase is not only involved in surviving ROS, but is absolutely essential for survival in the tick (13) and murine hosts. A potential mechanism for how this enzyme contributes to ROS detoxification is through the role of xanthine dehydrogenase in producing uric acid, which has been shown to be a potent antioxidant with strong free radical scavenging capabilities (23–25). As the bacterium is exposed to high levels of hypoxanthine (27) and ROS during the blood meal, we hypothesize that the bacteria may use *BB_*0554, *BB_*0555, and *BB_*0556 to shunt some portion of the excess hypoxanthine to generate uric acid and help neutralize the high levels of ROS. In line with this, we found that uric acid could rescue *BB_0554-BB_0556* ROS resiliency. Thus, a critical role for uric acid in *B. burgdorferi* pathogenesis appears likely.

This work ends with three important future directions. First, while the combination of our *in silico* analyses and our experiments using uric acid and allopurinol strongly suggest that the *BB_0554-BB_0556* operon is a xanthine dehydrogenase, we were unable to biochemically confirm the identity of the complex. We utilized a variety of methods but found uric acid to be in exceedingly low abundance inside *B. burgdorferi*. This made it infeasible for even high sensitivity mass spectrometry to reliably determine whether *ΔBB_0554-0556* bacteria had altered uric acid production. We hope that future work paired with technological advances will lead to a more concrete identity for this locus. Second, while our studies successfully link the operon to ROS resistance and to survival in ticks (13) and mice, we do not know whether the locus contributes to *in vivo* survival *through* ROS resistance. Interestingly, previous work demonstrated that *BB_0554, BB_0555*, and *BB_0556* transposon mutants are attenuated in wild-type and *gp91*^*phox*-/-^ mice (12), suggesting that Gp91-mediated superoxide production is not epistatic to *BB_0554-BB_0556*. However, this does not rule out that other constitutive sources of ROS in mammalian blood and tissues could mediate the survival defect of these mutants. Future studies will examine whether systematically neutralizing sources of ROS in animal systems relieves the pressure on *B. burgdorferi* and enables Δ*BB_0554-BB_0556* pathogenesis. Finally, while this work focused on ROS stress, reactive nitrogen species are also a critical stress that *B. burgdorferi* faces during transmission (2,3) and so understanding how the bacteria overcomes that pressure could improve our understanding of pathogenesis.

In sum, these findings increase our knowledge of *Borrelia burgdorferi* metabolism, ROS resistance, and pathogenesis. Understanding the mechanisms by which *B. burgdorferi* are able to resist ROS and other bloodborne insults could have important consequences on breaking the enzootic cycle. Conceptually similar approaches have already been tested, namely attempting to vaccinate white-footed mice against *B. burgdorferi* proteins expressed only in the tick (28). The logic for this study was that vaccination could result in serum-borne antibody and complement targeting *Borrelia* at the mammal-tick interface, killing the bacteria in both animal hosts. While attempts to leverage vaccines for this purpose have had varying effects depending on the model system, the potency of our findings here highlight a broader theme of potential anti-*Borrelia* therapeutics: the small genome and consequently limited metabolism encoded by the pathogen mean that disrupting any of the few known metabolic pathways could have potent effects due to reduced metabolic redundancy. Indeed the ability to specifically target *Borrelia* has been demonstrated by the identification of a spirochete-specific antibiotic (29). Improving our understanding of *B. burgdorferi* metabolism could therefore serve to greatly expand our arsenal of drugs to combat the pathogen.

## Materials and Methods

### Bacterial Strains and growth conditions

*B. burgdorferi* strains were grown in Barbour-Stoenner-Kelley II (BSK-II) medium in sealed tubes at 32°C with 1% CO_2_. *Escherichia coli* strains (Top10) for plasmid preparation were grown on Luria Bertani (LB) agar plates or in LB broth at 37°C. *E. coli* cultures contained either 50 μg/ml spectinomycin or 10 μg/ml gentamicin. The parental strain of the Tn library, the infectious *B. burgdorferi* strain 5A18NP1, was used as the wild-type strain in all studies and lacks two plasmids (lp56 and lp28-4)(30). The following antibiotics were used for selection in cultures of *B. burgdorferi* when appropriate: kanamycin at 200 μg/ml, gentamicin at 40 μg/ml, and streptomycin at 50 μg/ml. Tn mutants were obtained from the arrayed *B. burgdorferi* library (30). All individual Tn mutants used in this study were screened by polymerase chain reaction (PCR) at the locus of interest to confirm pure populations as previously described (12,31). All strains used in this study were routinely plasmid-typed to identify the loss of any plasmids required for murine or tick infection (32,33).

### Protein Modeling Prediction and phylogenetic tree

The secondary structure of each individual protein *BB_*0554, *BB_*0555, and *BB_*0556 were generated using Phyre2 (34). All three sequences were added as potential interacting partners and input into the SWISS-MODEL online software tools (35). These were visualized by the pyMOL program. The phylogenetic tree for each of the individual genes was generated using Phylogeny.fr (36). Following protein BLAST sequences were input into the “one click” function and a changes per site tree was generated by the PhyML Matrix program (36).

### RNA Extraction and RT-PCR Operon Analysis

5A18NP1 bacterial cells were grown to mid-logarithmic growth phase at 32°C. The culture was split in half with one set being used for genomic DNA extraction while the other was used for total RNA extraction using TRIzol (Invitrogen) following the manufacturer’s instructions. RNA samples were treated with the TURBO DNA-*free* kit (Invitrogen) to remove contaminating DNA. cDNA was prepared using random hexamers (Promega) and the ImProm-II Reverse Transcription System (Promega). Control reactions were performed in the absence of reverse transcriptase to control for the presence of genomic DNA. Sequences of the primers used to determine the identity of the operon are listed in Table 2. Following these procedures 2 μL of cDNA or control DNA was used as the template in PCRs using the pairs of primers listed in Table 2. PCR mixtures were incubated for 2 min at 95°C, subjected to 30 cycles of PCR, with 1 cycle consisting of 30 s at 95°C, 30 s at 53°C, and 45 s at 72°C, and finally incubated at 72°C for 5 min. PCR products were then resolved in 1% agarose gels and visualized by SYBR Safe (ThermoFisher) staining. Each RT-PCR experiment was replicated at least twice using different RNA preparations.

**Table 2:**
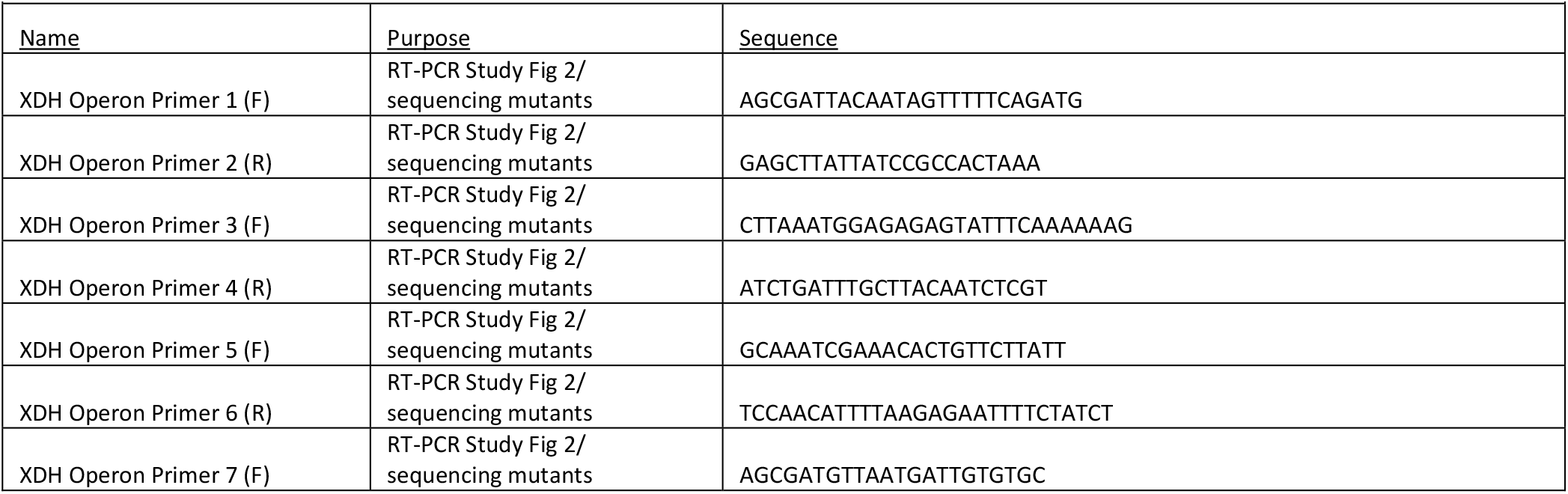
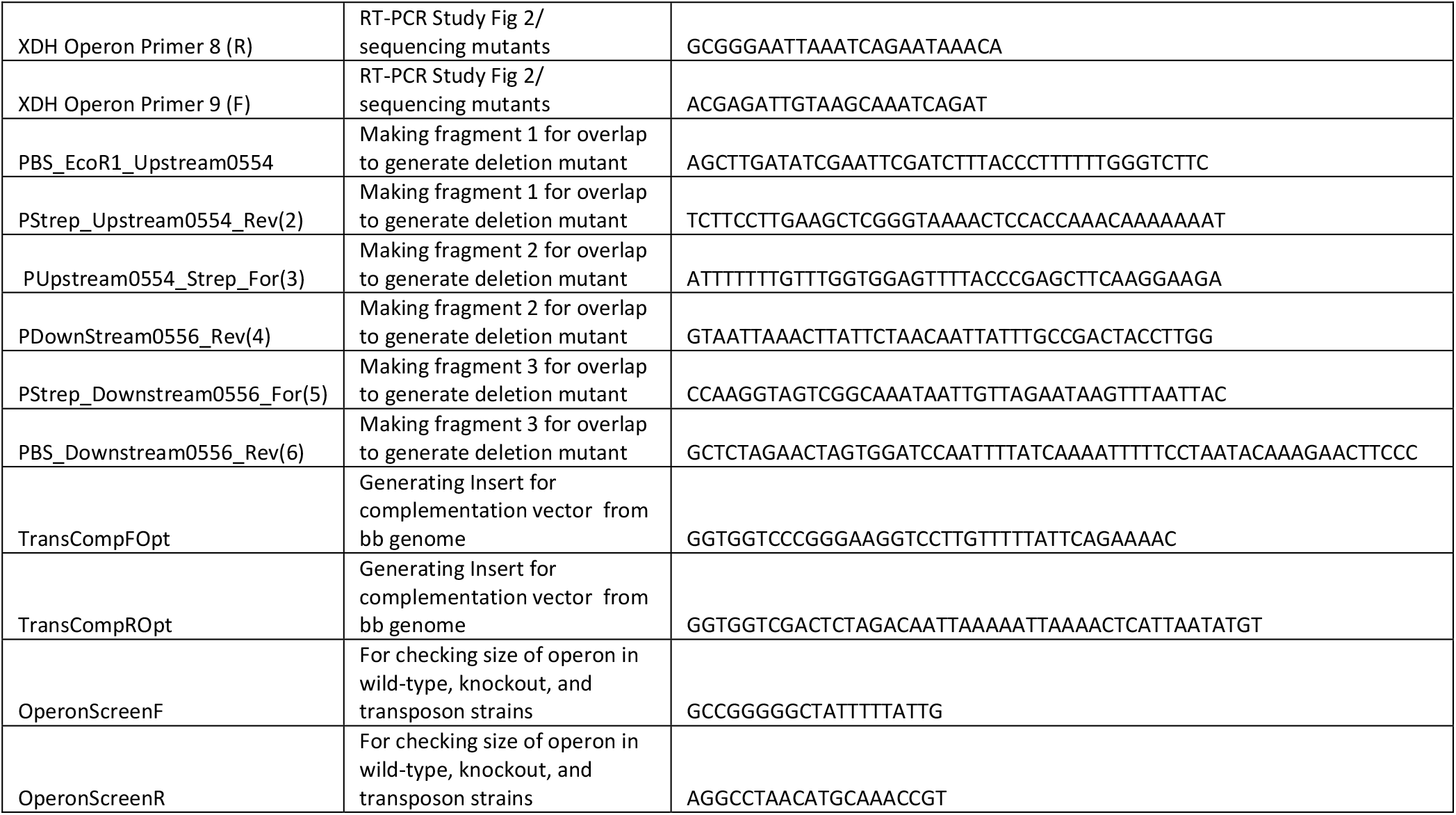
Oligonucleotides used in this study.

### Generation of *B. burgdorferi* mutant and complemented strains

Plasmid pJP01 was constructed to direct allelic exchange at the *BB_0554, BB_0555*, and *BB_0556* locus, resulting in the deletion of the operon’s open reading frame (Figure 2A). This plasmid was constructed by generating three PCR fragments : a 515 bp of sequence upstream of *BB_0554* (amplified from genomic DNA using primers PBS_EcoR1_Upstream0554 and PStrep_Upstream0554_R2, followed by a sequence containing the constitutive P*flgB* promoter and a streptomycin resistance gene (*aadA*, amplified from PKFSS1 using primers PUpstream0554_Strep_F3 and PDown0556_R4), followed by 524bp of sequence downstream of *BB_0556* (amplified from genomic DNA using primers PStrep_Downstream0556_F5 and PBS_Downstream0556_R6). The insert and pBlueScript plasmid were digested with EcoR1 and BamHI, which were subsequently ligated together using T4 DNA Ligase (New England Biolabs) and transformed into competent cells for selection. Prior to transformation into *Borrelia burgdorferi* the construct was linearized using KpNI and ECOR1.

Plasmid pJP02 was constructed for trans complementation of the *BB_0554, BB_0555, BB_0556* deletion mutant. This plasmid was generated by amplifying a DNA sequence containing all three genes as wells as the intergenic region upstream of *BB_0554* (273bp) from 5A18NP1 genomic DNA via PCR using primers TransCompFOpt and TransCompROpt. The resulting PCR products as well as Borrelia shuttle vector pBSV2-G were digested with XmaI and XbaI and the resulting PCR product and pBSV2-G backbone were ligated together using T4 DNA Ligase (New England Biolabs), followed by subsequent transformation into *E. coli*, generating PJP02 Plasmid pJP02 and the linearized deletion construct were introduced into *B. burgdorferi* by transformation as previously described (37,38). Potential transformants were confirmed by PCR with primers designed to detect either a replicating plasmid or a double crossover event, as appropriate, followed by dideoxy sequencing of the PCR product to confirm the expected nucleotide sequence.

### Determining *B. burgdorferi* sensitivity to ROS

The sensitivity of individual B. burgdorferi strains including the WT 5A18NP1, Tn:0556 mutant, the entire operon deletion as well as its complement to H_2_O_2_ was determined using an outgrowth assay similar to one previously described (39). Cells were grown for three days in BSK-II. The cell density of the cultures was determined using dark-field microscopy, and the cultures were diluted to a concentration of 2×10^7^/ml and grown overnight. Spirochetes were again counted under darkfield microscopy and 8.75×10^7^ cells were exposed to 0.125 μM or 0.185 μM H_2_O_2_ for four hours at 32°C. The concentration of H_2_O_2_ for each experiment was chosen to result in approximately 50% survival of the parental strain, which varied between bottles of H_2_O_2_. At the end of the stress exposure, the cells were diluted 5-fold in BSK-II. Cell density was quantified after three days, when the cell density of the untreated control reached ∼1×10^8^/ml. A percent outgrowth was determined for each strain by enumerating cell numbers using dark-field microscopy and then comparing the treated samples to the untreated controls. For experiments using allopurinol or uric acid, strains were exposed to the treatment (1mM allopurinol, 300nM uric acid) or DMSO control for two hours and washed twice prior to counting and exposure to H_2_O_2_.

### Subcutaneous Infection

Six-week-old male C57BL/6J mice were obtained from Jackson Laboratories and housed at the Tufts University Animal Facility for at least one week prior to infection. *B. burgdorferi* were grown for three days at 32°C and counted by darkfield microscopy. Spirochetes were resuspended in BSK-II at a density of 1×10^7^/mL and 200μL of this suspension (∼2×10^6^ total spirochetes) was injected subcutaneously in the posterior dorsal region of the mouse. After two weeks, ears, bladders, and ankles were harvested and transferred to 1mL BSK-II containing 50μg/mL rifampicin, 100μg/mL phosphomycin, and 5μg/mL amphotericin B. Cultures were monitored weekly for one month for spirochete growth by darkfield microscopy.

### Ethics Statement

Mice were maintained in the Tufts University Animal Facility. All experiments were performed following the guidelines of the American Veterinary Medical Association (AVMA) as well as the Guide for the Care and Use of Laboratory Animals of the National Institutes of Health. All procedures were performed with approval of the Tufts University Institutional Animal Care and Use Committee (IACUC, Protocol# B2018-98). Euthanasia was performed in accordance with guidelines provided by the AVMA and was approved by the Tufts University IACUC.

